# Microbiome dynamics associated with the infection of grey field slugs by the biocontrol nematode *Phasmarhabditis hermaphrodita*

**DOI:** 10.1101/2025.04.27.650856

**Authors:** Anh D. Ha, Dana K. Howe, Andrew J. Colton, Rory J. Mc Donnell, Dee Denver

**Affiliations:** Department of Integrative Biology, Oregon State University, Corvallis OR 97331, USA; Department of Crop and Soil Science, Oregon State University, Corvallis OR 97331, USA

## Abstract

The facultative-parasitic nematode *Phasmarhabditis hermaphrodita* has been used for many years as a biological control agent targeting slug pests. During the nematode’s infection process, the presence of certain bacteria has been suggested to affect the pathogenicity and efficiency of the nematodes in killing slugs, though the potential roles of different bacteria in affecting host-infection by nematodes remains unclear. In this study, we examined three experimental *P. hermaphrodita* populations cultured with three different bacteria: 1) *Escherichia coli* (EC), 2) a newly isolated *Pseudomonas sp.* Strain (PS) that co-cultured with a *P. hermaphrodita* strain found in Oregon, USA, and 3) the original complex bacterial community (BC) associated with the nematode. For each of the three treatments, we evaluated the pathogenicity of *P. hermaphrodita* at a low and high concentration towards adult *Deroceras reticulatum* (Mollusca: Gastropoda) and investigated the changes in the nematode microbiome structure before and after slug infection. Slugs exposed to EC low treatments (LT50: 8.61 days) and EC high treatments (LT50: 7.74 days) survived significantly longer than slugs exposed to PS high (LT 50: 5.68 days) and BC high (LT50: 5.92 days). Slugs in the BC low treatment (LT50: 6.77 days) survived significantly longer when compared to BC high, but survival was significantly shorter when compared to EC high. We identified a wide variety of taxa components (82 genera) in the community associated with the nematode pre-infection, most of which are of low abundance. In all bacterial treatments post-infection, the number of genera almost quadrupled and those taxa’s abundance changed remarkably, although the taxa with the highest abundance remained stable. We also observed four *Pseudomonas* amplicon sequence variants (ASVs) that increased in abundance after slug infection in the *Pseudomonas* treatment, which may suggest a role of the taxa in the infection process.

## Introduction

Invasive pest gastropod species such as the grey field slug, *Deroceras reticulatum*, are among the most widespread and damaging pests of agricultural and horticultural production systems, causing reductions in quality and yield loss on a broad range of crops including wheat, corn, legumes, vegetables, and fruits (Godan, 1983; Barker, 2002; Mc Donnell et al., 2009; Anderson et al., 2011). Slug pests have also been documented to vector several plant and human pathogens, such as *Alternaria brassicicola* (causing black spot disease in brassica), *Escherichia coli* (causing food poisoning) (Mc Donnell et al., 2009), and *Angiostrongylus cantonensis*, the causal agent of the potentially lethal eosinophilic meningitis (Centers for Disease Control and Prevention, 2023).

Conventional slug control methods primarily rely on chemical solutions, using active molluscicide ingredients such as metaldehyde, iron phosphate, and carbamates (Barker, 2002; Mc Donnell and Anderson, 2017). However, environmental concerns exist regarding the potential for these chemical molluscicides to harm non-target organisms (Aktar et al., 2009; UK Department for Environment, 2020; Tudi et al., 2021).

Furthermore, there is evidence that target slugs might be developing resistance to these molluscicide compounds (Adomaitis et al., 2022). Therefore, alternative slug control strategies, such as biological control approaches, are increasingly common in integrated pest management systems.

*Phasmarhabditis hermaphrodita* is a facultatively parasitic, rhabditid nematode species that infects a variety of slug and snail species. The potential of *P. hermaphrodita* for biocontrol against slugs was realized as early as 1988, when it was confirmed to be highly virulent against a wide range of pest gastropods (Wilson et al., 1993; Rae et al., 2007). The nematode has been sold commercially under the trade name Nemaslug® in Europe for over 20 years; still, it is not currently available for commercial purchase in the United States, due to regulatory concerns regarding incomplete information about the effects of this nematode on North America-native gastropod species. Recent work by our research team demonstrated that *P. hermaphrodita* infects and kills *Monadenia fidelis*, a non-target snail species endemic to the Pacific Northwest of North America (Denver et al. 2024). Discoveries of *P. hermaphrodita* in California (De Ley et al., 2014) and subsequently in Oregon (Mc Donnell et al., 2018; Howe et al., 2020) motivated further exploration of this nematode as a potential biocontrol agent for slugs in the United States.

Regarding the mechanism of infection, *P. hermaphrodita* shares some similarities with the well-studied entomopathogenic nematodes (EPNs) of the genera *Steinernema* and *Heterorhabditis*, which have been commonly used as biological control agents of insect herbivores (Lacey and Georgis, 2012). The infective juvenile-stage *P. hermaphrodita* lives freely in the soil and gains entrance to the hosts upon exposure, usually through the mantle cavity. Once inside, infective juveniles develop into self-fertilizing hermaphroditic adults, reproduce, and the infection spreads to the entire body of the slug host, eventually causing its death within 4-21 days. The nematodes continue to consume the slug carcass until the food source is depleted, and their new infective juveniles again move into soil to search for new susceptible hosts (Tan and Grewal, 2001a; Rae et al., 2007).

Despite sharing many similarities in the life cycle with the EPNs of the genera *Steinernema* and *Heterorhabditis*, which respectively engage in an obligate, specific mutualisms with *Xenorhabdus* and *Photorhabdus* bacteria (Forst et al. 1997; Sicard et al., 2004), *P. hermaphrodita* has a more uncertain and understudied relationship with its bacterial associates. The scientific literature focusing on the role of bacteria in the lifestyle, virulence, and efficacy of *P. hermaphrodita* presents inconsistent findings. In searching for a bacterium suitable for industrial-scale production of *P. hermaphrodita* as a commercial bio-molluscicide, Wilson and colleagues selected *Moraxella osloensis* (Wilson et al., 1995b, 1995c), which was later employed in the widely used Nemaslug® product. However, they also observed that *P. hermaphrodita* could feed on and carry out slug infection with many other bacterial partners, including *Pseudomonas fluorescens* and *Pseudomonas paucimobilis.* Direct injection of *M. osloensis* into slugs did not cause significant mortality in the mollusk hosts (Wilson et al., 1995a, 1995b). On the contrary, Tan and Grewal proposed that the sole agent responsible for pathogenicity was *M. osloensis* by producing endotoxin, and *P. hermaphrodita* nematode served merely as a vector that transports the virulent bacterium into the shell cavity of the slug (Tan and Grewal, 2001b, 2002). Meanwhile, other studies suggested that *P. hermaphrodita* lacked a specific obligate partner in killing the slugs, and bacteria did not have a discernible influence on virulence (Rae et al., 2010; Sheehy et al., 2022). The precise roles of nematodes and bacteria in slug infection remain unclear, hindering the optimization of biocontrol methods using the nematode.

To further investigate the relationships between nematodes and bacterial partners during slug infections, we conducted a set of slug infection trials in an effort to evaluate the pathogenicity of *P. hermaphrodita* nematodes reared on:

1. *Escherichia coli* strain OP50, a known non-pathogenic bacterium, usually used as the common food source for laboratory populations of the famous model organism *Caenorhabditis elegans*.
2. One *Pseudomonas* sp. strain isolated from the microbial community associated with a *P. hermaphrodita* nematode found in Oregon.
3. The complex bacterial community that co-cultured with *P. hermaphrodita* found in Oregon.

Our results contribute insights into the dynamics of microbiomes during nematode-slug infection processes, and further evidence that *P. hermaphrodita* can infect and kill hosts with a variety of bacterial associates.

## Materials and Methods

### Culturing of nematodes and bacteria

The *P. hermaphrodita* strain we used in this study (DL 309) was isolated from a dead slug collected in Oregon (Howe et al. 2020). The nematodes were maintained in the lab using on standard nutrient growth media (NGM) agar plates with co-cultured bacteria as a food source. NGM plates were prepared using autoclave-sterilized media, equipment, and sterile plastic plates (VWR International, Lutterworth, UK) and seeded with the appropriate bacterial cultures. Prior to our study, nematodes were bleached with sodium hypochlorite 0.25M solution following standard worm bleaching protocol (Stiernagle, 2006) to minimize bacterial carry-over from previous culturing conditions. The nematodes were monitored on plain NGM agar plates for 24 hours to ensure no bacterial growth, then harvested and washed once with sterile M9 buffer and twice with molecular water. Next, nematodes were transferred onto fresh NGM plates (Stiernagle, 2006) seeded with one of the three designated bacterial food sources: *E. coli* OP50 (EC)*, Pseudomonas sp.* (PS)., and the original bacterial community (BC) associated with the nematodes. The *E. coli* OP50 culture was obtained from the *Caenorhabditis* Genetics Center at the University of Minnesota. The *Pseudomonas* strain was isolated from the original bacterial community that co-cultured with a *P. hermaphrodita* sampled at the Oregon State University campus (Mc Donnell et al., 2018). We confirmed the genus identity of the strain with 16S rRNA and *rpoB* gene sequencing.

### Infection Assay Procedure

#### Experimental setup

Grey field slugs (*D. reticulatum*) were collected from several ryegrass seed production fields around Tangent, Linn County, Oregon one day prior to the experiment and maintained in a growth chamber (Thermo Scientific Precision Model 818) at 18℃ and 12-h photoperiod. The infection assay was performed in 16 oz plastic round containers (13.8 cm in height x 11cm in diameter), each containing 25g of sterilized damp EarthGro® topsoil. Soil was dampened in each container by adding 10 ml of deionized water and mixing thoroughly. The containers were stored in the same growth chamber.

Our infectivity assay followed the approach described by Mc Donnell et al. 2020. Immediately before the trial, nematodes were harvested from NGM plates and washed once with sterile M9 buffer and twice with molecular water. The number of nematodes were estimated by counting 1 mL sub-samples and then resuspended in 5 mL of sterile water and distributed into 10 replicates per nematode-bacteria combination, five with a concentration of ∼8,000 nematodes/ml (‘high dose’) and five with a concentration of ∼4,000 nematodes/ml(‘low dose’). Nematodes were pipetted onto the soil to the final concentration of ∼210 worms/cm² for low dose, or ∼420 worms/cm² for high dose. In total, 30 containers were set up [3 combinations x 2 doses x 5 replicates = 30]. Six healthy, mature adult *D. reticulatum* slugs (>100 mg) were added to each of the 30 containers. In our previous pilot studies, no sign of aggression was documented with this number of slugs cohabiting in the same container. Five additional containers, each holding six slugs and no nematodes were set up as negative controls. All slugs were fed with sterilized carrots during the course of the experiment.

The slugs were monitored daily for symptoms of nematode infection (e.g. swelling of the mantle, emaciation of the slug body, exposure of the internal shell, and nematodes visible on the slug body) and mortality for 15 days. Lethal Time 50 (LT50) and fiducial limits (95%) were calculated for the slug survival data using probit analysis. Abbott’s formula was used to correct for control mortality. Statistically significant differences (P<0.05) exist when there is no overlap between the 95% fiducial limits. All analyses were carried out using IBM® SPSS® Version 24.

#### Collection of nematode-bacteria samples before and after infection

For each of the three bacteria examined in this study (EC, PS, BC), three biological replicates were collected, both before and after host slug infection. Prior to infection, we collected nematode samples in a fixed volume of 200 µL nematode suspension in sterile M9 solution on a semi-sterile lab benchtop as described above. Immediately before the infection assay, three replicates of nematode suspensions at high concentration (8,000 nematodes/ml) were taken out from each of the three combinations, totaling nine samples. The worms were washed once with M9 buffer, and then twice with distilled water. We also included twelve negative controls to assess the contamination introduced during the preparation procedure and sequencing: three samples of the M9 buffer, three samples of the distilled water used during the infection assay step, three samples of distilled water used during the nematode washing step, and three DNA Extraction kit blanks. The nine pre-infection nematode samples (hereafter referred to as “PreInf” sample) and twelve negative controls were stored at −80 °C immediately after collecting.

After host slug infection, nematodes and bacteria from the high-dose nematode treatment were collected in three biological replicates. For each of the three bacteria treatments, we collected three slug carcasses, each from a separate container, to extract their nematode populations. Immediately following the death of a slug, the carcass was submerged in a sterile M9 solution to elute nematodes. Collected nematodes were washed once with M9 buffer and then twice with molecular biology grade water (VWR International, Lutterworth, UK). The nematode samples (hereafter referred to as “PostInf” samples) were then stored at −80°C until DNA extraction. The simplified naming scheme for the six different bacterial treatment types used hereinafter is summarized in Table S1.

#### DNA Isolation, 16S rRNA Amplification and Sequencing

Prior to DNA isolation, nematode samples were homogenized thoroughly by bead beating. We extracted DNA from the homogenate and from the negative control samples using the PowerSoil DNA Extraction kit (MOBIO, Carlsbad, CA, USA). DNA extractions were then quantified using a fluorescent plate reader at the OSU Center for Genome Research and Biocomputing (CGRB).

16S library preparation, amplification, and sequencing were conducted by the OSU CGRB. Amplicon libraries were prepared following the standard protocol by Illumina (16S Metagenomic Sequencing Library Preparation, 2013). Primers targeting the V3-V4 region (forward: 5’TCGTCGGCAGCGTCAGATGTGTATAAGAGACAGCCTACGGGNGGCWGCAG3’; reverse: 5’GTCTCGTGGGCTCGGAGATGTGTATAAGAGACAGGACTACHVGGGTATCTAATCC 3’) were used for PCR amplification (Klindworth et al., 2013). Subsequently, the PCR products were processed through clean-up, index PCR, library quantification, normalization, and pooling. Libraries were sequenced using paired-end MiSeq v2 Nano 300 bp platform (Illumina, USA).

#### DNA Sequence Data Processing

We used the DADA2 pipeline (Callahan et al., 2016) to trim reads, merge paired reads, denoise, quality filter, and infer amplicon sequence variants (ASVs). ASVs represent exact 16S rRNA gene sequence variants resolved to the level of single-nucleotide dissimilarity. Compared to the commonly used operational taxonomic units (OTUs) with a threshold of nucleotide difference of 97%, ASVs have been shown to offer better reproducibility, reusability, and comprehensiveness in 16S microbiome analyses (Callahan et al., 2017).

Phylogenetic analyses and representations of the ASVs were constructed using MAFFT alignment (Katoh et al., 2002) and FastTree 2 (Price et al., 2010). To assign taxonomy to the ASVs, we utilized a self-trained classifier, trained on the Silva Project’s database (release 132) (Quast et al., 2013). Putative contaminant sequences were identified and filtered out of sample data using the R package decontam (Davis et al., 2018). ASVs were considered a contaminant if they were more prevalent in negative controls than in positive samples and/or their frequency significantly varied inversely with sample DNA concentration (probability threshold p < 0.1).

We utilized the Phylum, Genus, and ASV levels as units for subsequent taxa analyses. Since different taxonomic classifiers trained on different databases may differ at lower taxonomic ranks (Balvočiūtė and Huson, 2017), the Phylum level was chosen as the main unit for taxa analysis; however, the Genus and ASV level were also selected for a finer resolution and subsequent composition analysis.

#### α-Diversity Analyses

We used four metrics: Chao1, Shannon Index, Inverse Simpson, and Faith’s Phylogenetic Diversity (PD) to compare and evaluate α-diversity from different approaches. Chao1 index is an estimator of species richness i.e. the expected total number of OTUs/ASVs in the sample given all the species were identified. It weighs the low abundance taxa (i.e. only singletons and doubletons) to infer the number of missing species (Chao, 1984). The Shannon Index estimates the overall richness and also the evenness/uniformity between the taxa present in the sample (Spellerberg and Fedor, 2003). Also focusing on the species richness and evenness, however, the Inverse Simpson index emphasizes more on species evenness (Simpson, 1949), while the Shannon Index gives greater weight to the species richness. Faith’s PD metric considers the phylogeny of taxa to estimate diversity and is proportional to how much of the OTU/ASV phylogenetic tree is covered by the taxa present (Faith, 1992).

To calculate the Shannon Index and Inverse Simpson, the ASV table was rarefied ten times to a depth of 100,000 reads to lessen the random biases introduced by random subsampling. Rarefying the ASV table to a fixed number of reads per sample is necessary for calculations of the Shannon Index and Inverse Simpson, as these two abundance-based indices could be substantially affected by the differences in the total number of reads between samples. The two indices were calculated for each of the ten rarefied tables using the package phyloseq (McMurdie and Holmes, 2013), and the mean values were used for α-diversity analysis. As for Chao1, which depends on low-abundance taxa, the ASV table was not rarefied to retain all ‘rare’ taxa. The Faith’s PD index was calculated using the package picante (Kembel et al., 2010). We assess the significance of the differences among the bacteria-nematode combinations at the same time point using Kruskal-Wallis tests. Mann-Whitney U tests or paired t-tests were applied, as appropriate, to test for the difference between the two time points before and after infection. The normality of diversity values was assessed using the Shapiro-Wilk normality test.

#### β-Diversity Analyses

To assess the difference in the microbial community between samples, we used four β-diversity metrics: two abundance-weighted β-diversity indices (Bray-Curtis and weighted UniFrac) and two presence-absence indices (Jaccard and unweighted UniFrac). While Bray-Curtis and Jaccard are non-phylogenetic metrics, weighted and unweighted UniFrac also take into account the phylogeny of the bacterial community composition to assess differences between samples. Specifically, UniFrac distances are based on the sum of branch length shared between samples on the phylogenetic tree constructed from all the 16S rRNA sequences from all communities of interest. To test whether the groups differ significantly from each other, we conducted the permutational multivariate analysis of variance (PERMANOVA) test using the package vegan (Oksanen et al., 2012).

## Results

### Pathogenicity and efficacy of P. hermaphrodita in killing slugs

Slugs that were exposed to the PS high rate treatment (LT50: 5.68 days) survived for significantly (P<0.05) fewer days than those treated with both the EC high (LT50: 7.74 days) and low (LT50: 8.61 days) rates (Figure 1). For BC high (LT50: 5.92 days), slugs survived significantly (P<0.05) fewer days than BC low (LT50: 6.77 days) and both EC rates. Lastly, slugs exposed to BC low survived significantly fewer days than those treated with EC high (Figure 1).

**Figure 1.**
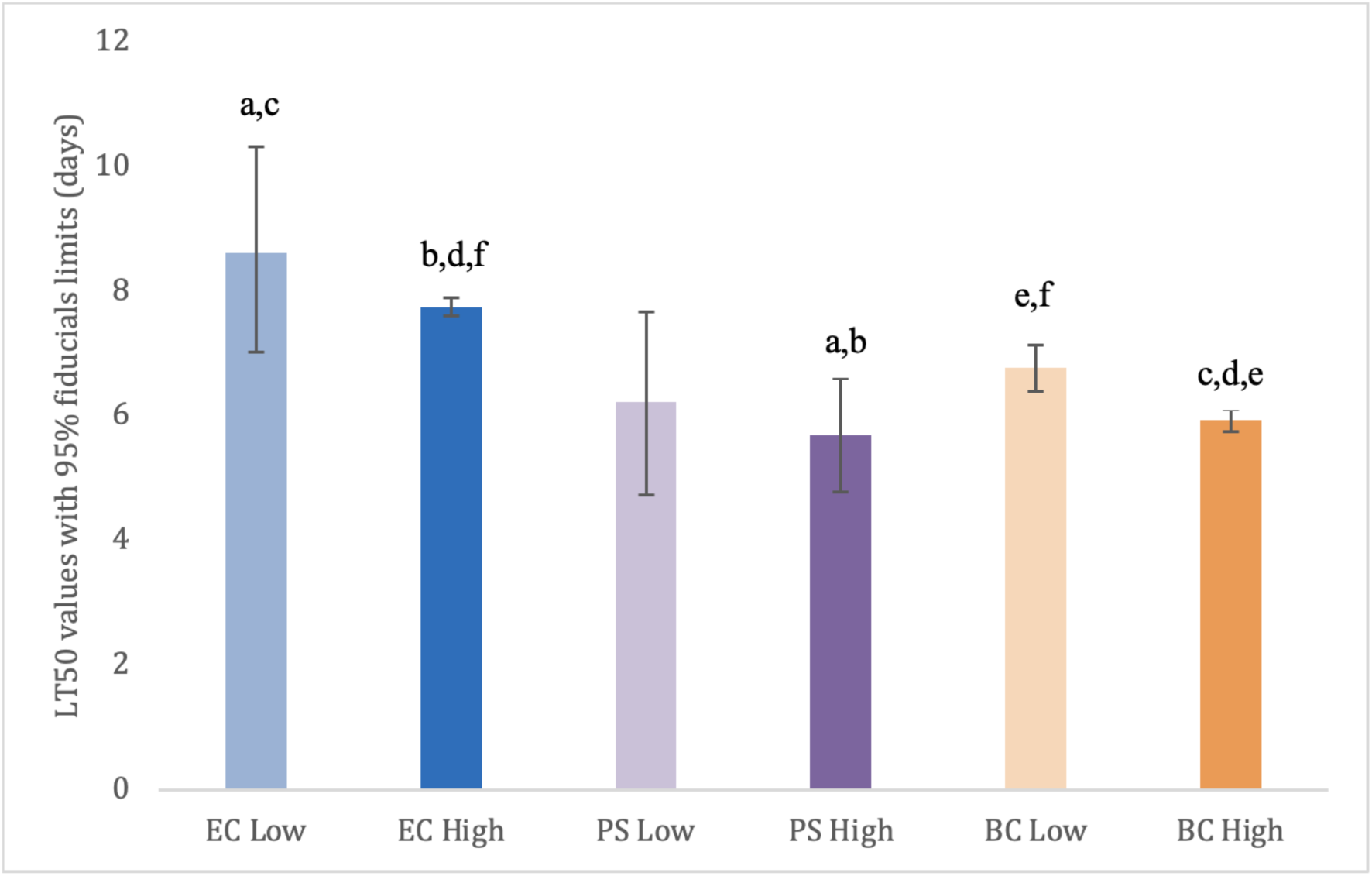
Lethal time 50 (days) with 95% fiducial limits for adult *D. reticulatum* when exposed to low and high rate treatments of *P. hermaphrodita.* Nematodes were paired with one of three designated bacterial food sources: *E. coli* OP50 (EC)*, Pseudomonas sp.* (PS), and the original bacterial community (BC) associated with the nematodes. Treatment data were corrected for control mortality using Abbott’s formula. Bars with the same lowercase letter indicate a statistically significant difference (P<0.05) between treatments. These difference exists when there is no overlap between the fiducial limits. Lethal time 50 values (95% fiducial limits): EC low rate = 8.61 (7.01 – 10.31), EC high rate = 7.74 (7.60 – 7.88), PS low rate = 6.23 (4.73 – 7.66), PS high rate = 5.68 (4.78 – 6.59), BC low rate = 6.77 (6.39 – 7.13), BC high rate = 5.92 (5.75 – 6.09).

### Bacterial 16S rRNA Sequence Data

After quality filtering and removing chimeric sequences, we obtained a total of 10,485,882 reads, with an average of 582,549 reads and a median depth of 349,960 reads per sample. 2,938 ASVs (i.e. taxa) were identified. The decontamination procedure removed 131 ASVs, leaving 2,807 unique ASVs for subsequent analyses. The components of the removed contaminants are shown in Figure S1. The filtered data set then contained 10,422,195 reads in total, with an average depth of 579,010 reads and a median 343,942 reads per sample. Details of the number of quality-controlled reads obtained in each sample, as well as the total number of reads, median, and mean values of reads per nematode-bacterial combination were listed in Table S2.

### Microbiome shifts before and after slug infection experiment

The number of different genera detected in each sample included in the analysis is listed in Table 1. On the Phylum level, the majority of bacterial components in all samples were identified as members of the phyla Bacteroidetes and Proteobacteria. We observed a shift in composition towards Proteobacteria in the BC-PostInf and PS-PostInf samples compared to their PreInf counterparts, while in the EC-PostInf samples, there was a shift towards Bacteroidetes (Fig. 2A). In general, the number of genera in all samples increased almost fourfold after infection. The abundance of the main genera also altered remarkably between PreInf and PostInf samples (Fig. 2B).

**Figure 2.**
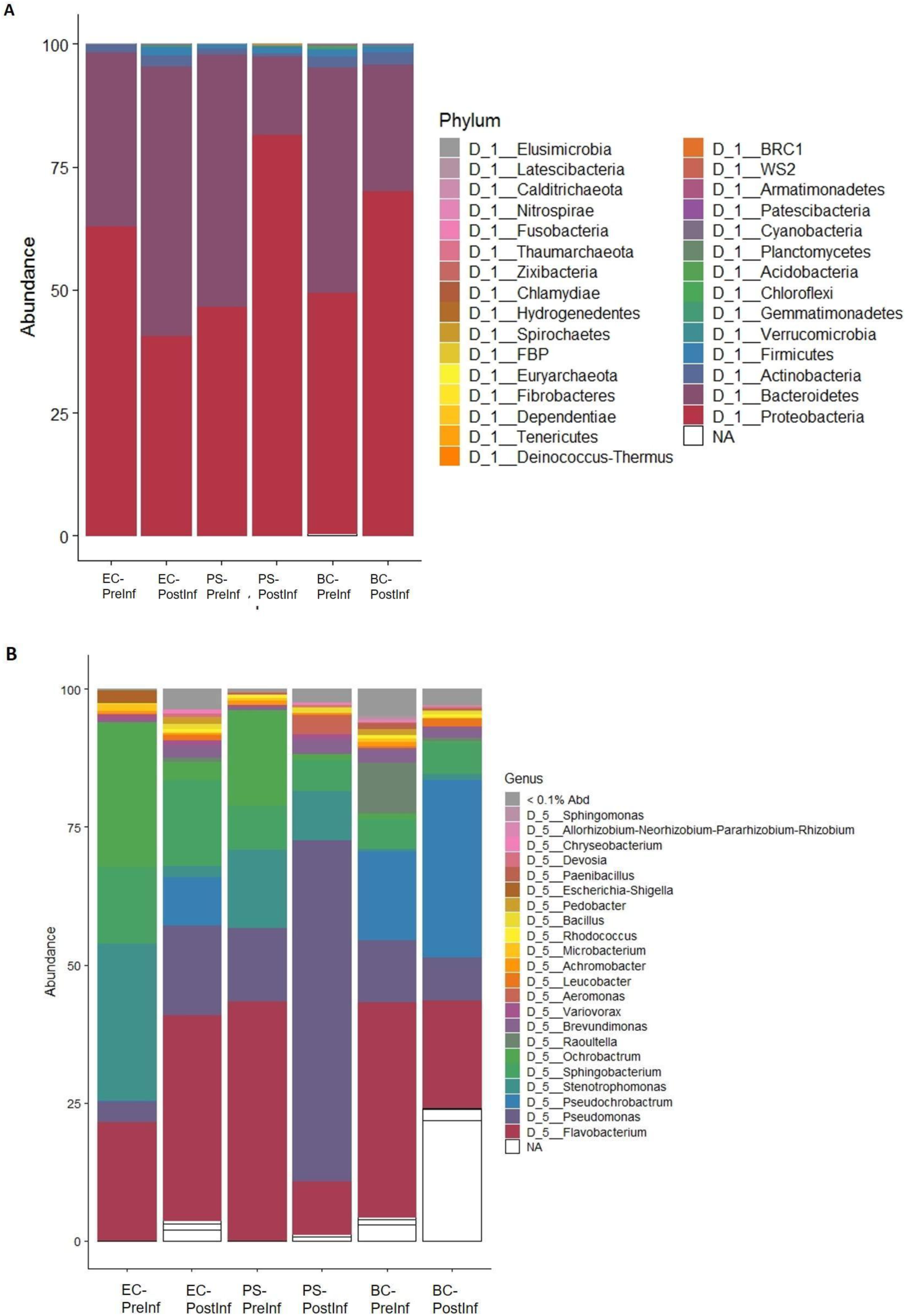
Composition of bacterial communities associated with *P. hermaphrodita* on the (A) Phylum and (B) Genus level.

**Table 1.** The number of different genera detected in each sample.

| **Bacterial treatment** | **Sample (before infection)** | **No. of genera** | **Sample (after infection)** | **No. of genera** |
| --- | --- | --- | --- | --- |
| *E. coli* | EC-PreInf-1 | 30 | EC-PostInf-1 | 108 |
|  | EC-PreInf-2 | 31 | EC-PostInf-2 | 185 |
|  | EC-PreInf-3 | 32 | EC-PostInf-3 | 80 |
| *Pseudomonas sp.* | PS-PreInf-1 | 22 | PS-PostInf-1 | 108 |
|  | PS-PreInf-2 | 27 | PS-PostInf-2 | 115 |
|  | PS-PreInf-3 | 24 | PS-PostInf-3 | 107 |
| Bacterial complex | BC-PreInf-1 | 66 | BC-PostInf-1 | 130 |
|  | BC-PreInf-2 | 36 | BC-PostInf-2 | 182 |
|  | BC-PreInf-3 | 35 | BC-PostInf-3 | 144 |

In *P. hermaphrodita*’s original microbiome (i.e. the BC-PreInf community), 15 phyla were identified with Bacteroidetes being the most abundant phylum, followed by Proteobacteria. On the Genus level, the most abundant groups were *Pseudochrobactrum, Flavobacterium, Raoultella,* and *Pseudomonas* out of the 82 genera observed (Fig. 2B). Details of relative taxa abundance on the Phylum and Genus level in the BC-PreInf communities were shown in Table S3A.

The numbers of ASVs unique to specific sample sets and shared among sample sets were illustrated in Fig. 3. In general, the number of ASVs unique to a single sample set was higher than those shared between different sets. The BC-PostInf samples had the largest number of unique ASVs (i.e. 600) that were not shared with any other type of samples, followed by the EC-PostInf and PS-PostInf (451 and 337 ASVs, respectively). Comparing PreInf and PostInf samples, the BC-PreInf and BC-PostInf communities shared 82 common ASVs, while PS-PreInf and PS-PostInf shared 36 ASVs, and 44 ASVs were shared between EC-PreInf and EC-PostInf samples.

**Figure 3.**
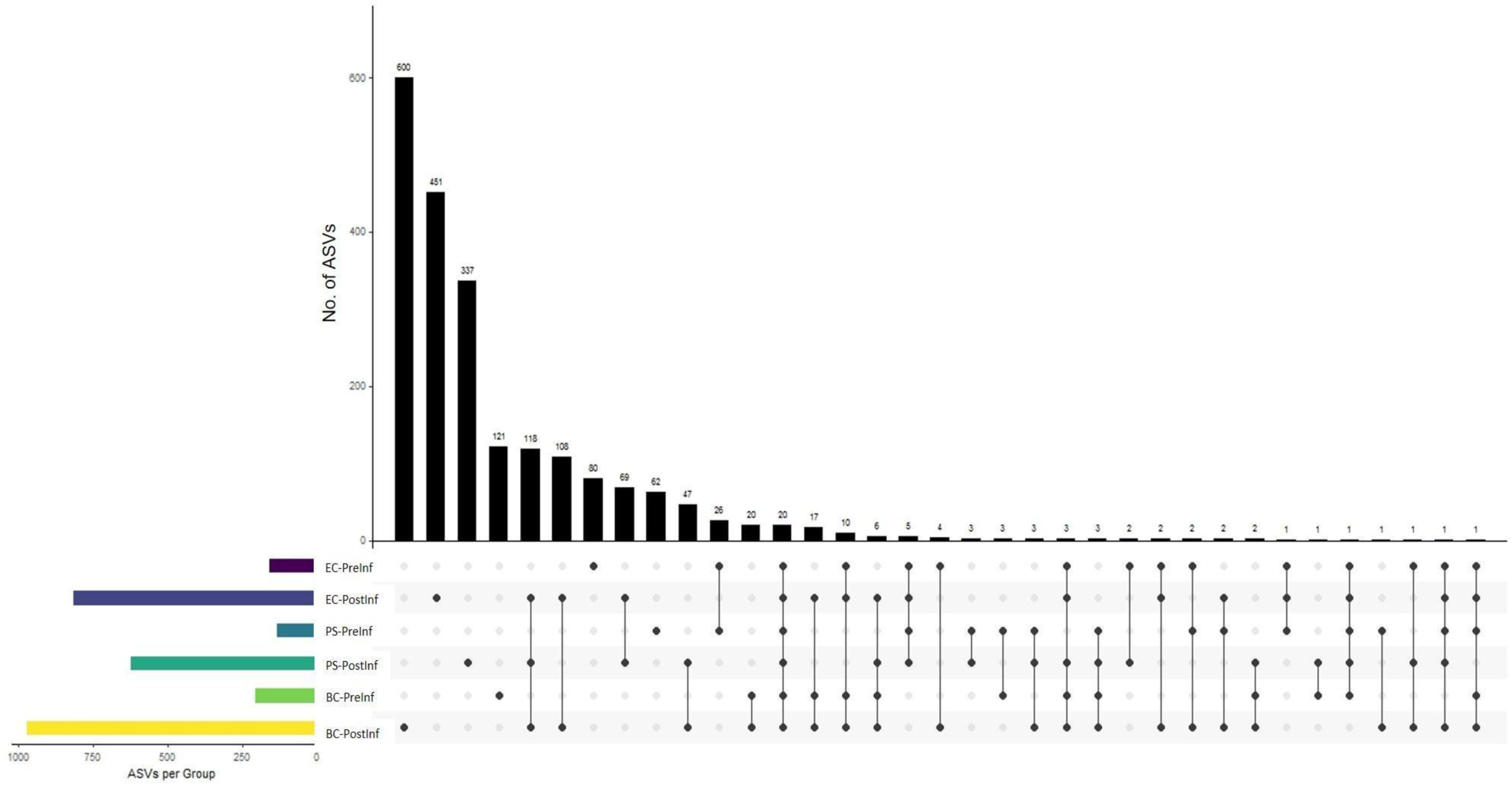
The ASV components present in each of the sample sets. Each horizontal colored bar on the left indicates the number of unique ASVs per sample group. Each vertical bar in the plot illustrates the number of ASVs detected in the samples. The dots (bottom) correspond to the presence of those numbers of ASV in the sample sets, and co-appearance of one dot in multiple samples demonstrates the number of ASVs that are shared among those different sets.

### Patterns of change in α-diversity

We evaluated α-diversity using the four metrics Chao1, Faith’s PD, Inverse Simpson, and Shannon Index (Fig. 4A). Chao1 index had a significantly higher median across all PostInf samples compared to PreInf (Mann-Whitney U test, p = 4.114e-05). There was suggestive evidence that Faith’s PD and Shannon indices were lower in PreInf-than PostInf-samples (Mann-Whitney U, p = 0.06 and p = 0.09, respectively).

**Figure 4.**
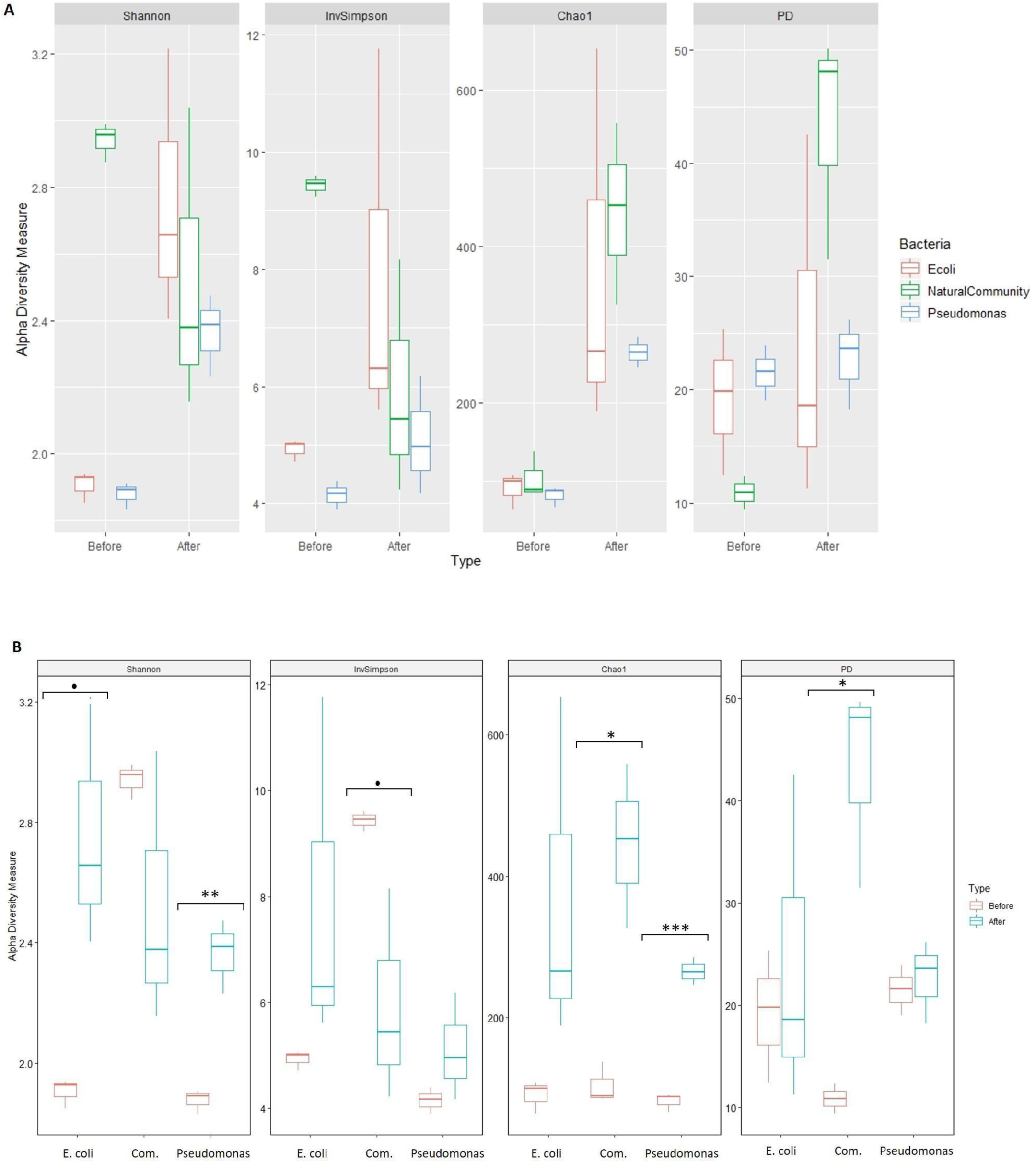
α-diversity indices Shannon, Inverse Simpson, Chao1 and Faith’s Diversity of samples grouped by (A) Time of collection and (B) Bacterial treatment. Significant differences of the metrics between pre- and post-infection of each bacterial treatment are indicated with the significant code: (***) 0-0.001, (**) 0.001-0.01, (*) 0.01-0.05, (.) 0.05-0.1, () > 0.1.

Analysis of the EC microbiomes suggested some evidence that the mean Shannon index increased after the slug infection process (Welch two-sample t-test, p = 0.068). The BC samples showed a moderately significant increase in Faith’s PD and Chao1 value (p = 0.029 and p = 0.03) and a suggestive decrease of the Inverse Simpson index. The PS-PostInf samples demonstrated strong evidence of an increase in Shannon and chao1 index (p = 0.01 and p < 0.0004, respectively) compared to PS-PreInf. The changes in α-diversity of each bacteria-nematode combination at the two time points were summarized in Fig. 4B and detailed in Table 2.

**Table 2.** Changes in α-diversity indices of each bacteria-nematode combination after infection trial.

| **Diversity metrics** | ***E. coli* (EC)** | **Bacterial Complex (BC)** | ***Pseudomonas sp.* (PS)** |
| --- | --- | --- | --- |
| Shannon | **Increase** (p = 0.068) | No significant difference (p = 0.25) | **Increase** (p = 0.01) |
| Faith's PD | No significant difference (p = 0.67) | **Increase** (p = 0.029) | No significant difference (p = 0.7) |
| Inverse Simpson | No significant difference (p = 0.27) | **Decrease** (p = 0.09) | No significant difference (p = 0.24) |
| Chao1 | No significant difference (p = 0.19) | **Increase** (p = 0.03) | **Increase** (p < 0.0004) |

There was some evidence that the diversity in all bacteria-nematode combinations prior to infection was not equal (Kruskal-Wallis, p = 0.05). After the infection assay, α-diversity indices between the three bacteria-nematode combinations did not differ significantly (Kruskal-Wallis test, p = 0.3012).

### Patterns of change in β-diversity

We applied four β-diversity indices to evaluate the dissimilarity between samples of the three combinations sampled before and after the infection assay: Bray-Curtis dissimilarity, Jaccard, unweighted, and weighted UniFrac. Bray-Curtis results showed a clear pattern of distinct sample clustering (Fig. 5). The bacterial community structures were significantly different between the EC, PS, and BC sample sets (PERMANOVA, p = 0.001, R2 = 0.541). This result was robust to Jaccard and Weighted UniFrac indices (PERMANOVA, p = 0.001 for both), but not to UniFrac (PERMANOVA, p = 0.06). Between samples collected PreInf and PostInf, Bray-Curtis dissimilarity, Jaccard, and UniFrac indices indicated a significant difference (PERMANOVA, p = 0.02, p = 0.01, p = 0.001, respectively); however, Weighted UniFrac did not show a clear deviation (p = 0.25).

**Figure 5.**
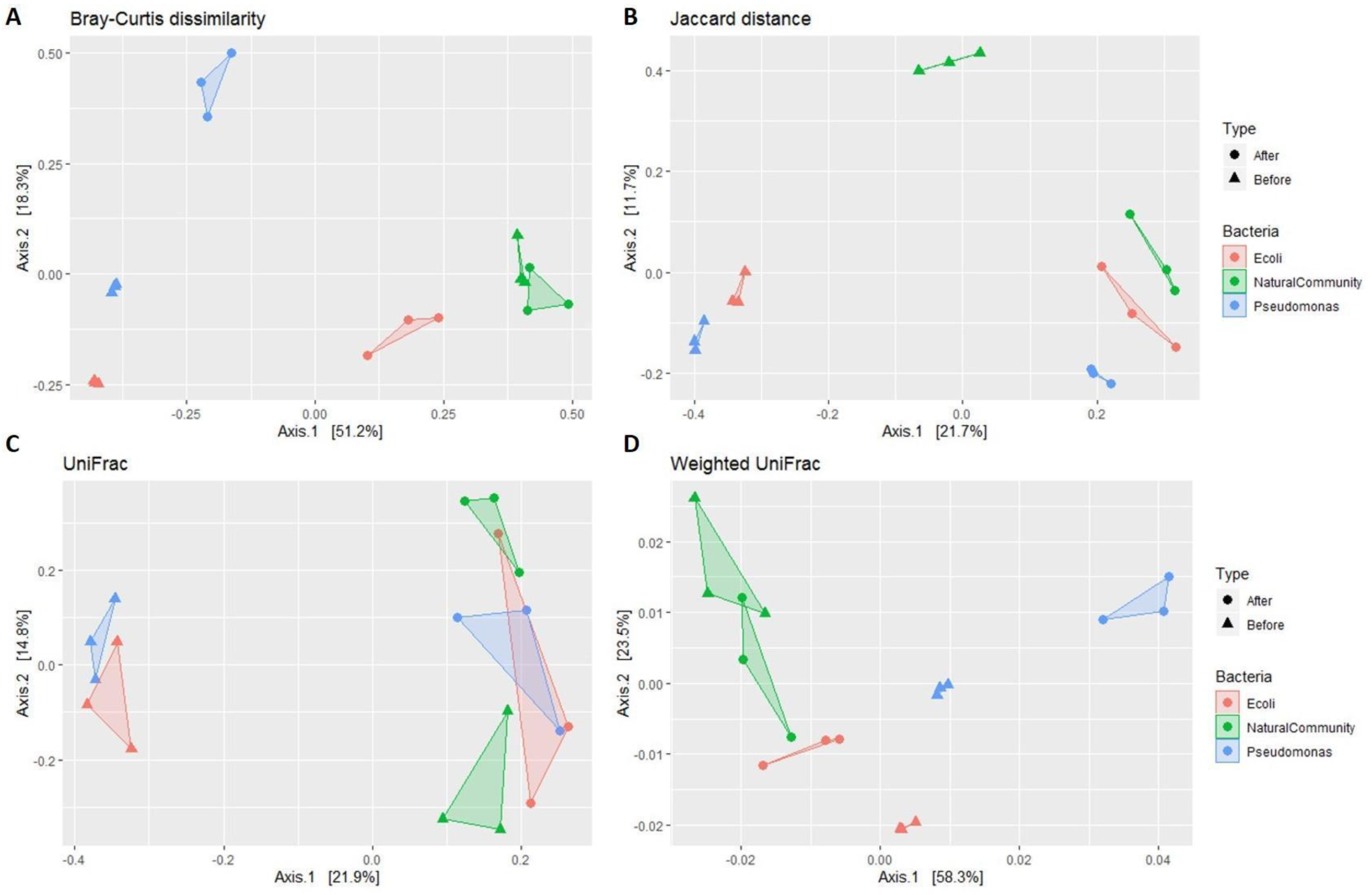
β-diversity indices of the microbiome samples. Principal coordinate analysis (PCoA) with a (A) Bray-Curtis dissimilarity, (B) Jaccard distance, (C) Unweighted UniFrac distance, and (D) Weighted UniFrac distance. The samples are colored by the type of bacterial treatment and shaped by the time of collection.

### *Pseudomonas* ASV expansion after slug infection in the PS samples

We analyzed the ASV components of the PS microbiome, with specific focus on those deriving from *Pseudomonas* spp., to evaluate the potential effects of the infection process on this group of bacteria (Fig. 6). The microbiome of PS-PostInf only shared a total of 36 ASVs with PS-PreInf, four of which were identified to be of the genus *Pseudomonas.* The relative abundance of these four ASVs in PS-PreInf samples was 13.23% of the total, and expanded to 59.58% in PS-PostInf samples. Out of these four ASVs, one was present in all three bacterial-nematode combinations sampled both before and after infection, and displayed an increase of 7.5%, 3.6%, and 3.2% in EC, PS, and BC communities, respectively. The three other *Pseudomonas* ASVs were only detected in the microbiomes of PS and BC nematodes, both before and after the infection assay; they were absent in all EC samples. While their abundances all increased after the course of infection in PS samples - by 6.9%, 30.5%, and 5.5%, there was a consistent decrease of these ASVs by 1.38%, 4.13%, and 0.73% from BC-PreInf to BC-PostInf samples. Two *Pseudomonas* ASVs only present in PS-PreInf accounted for just 0.004% abundance;14 ASVs unique to the PS-PostInf had a total relative abundance of 1.11%.

**Figure 6.**
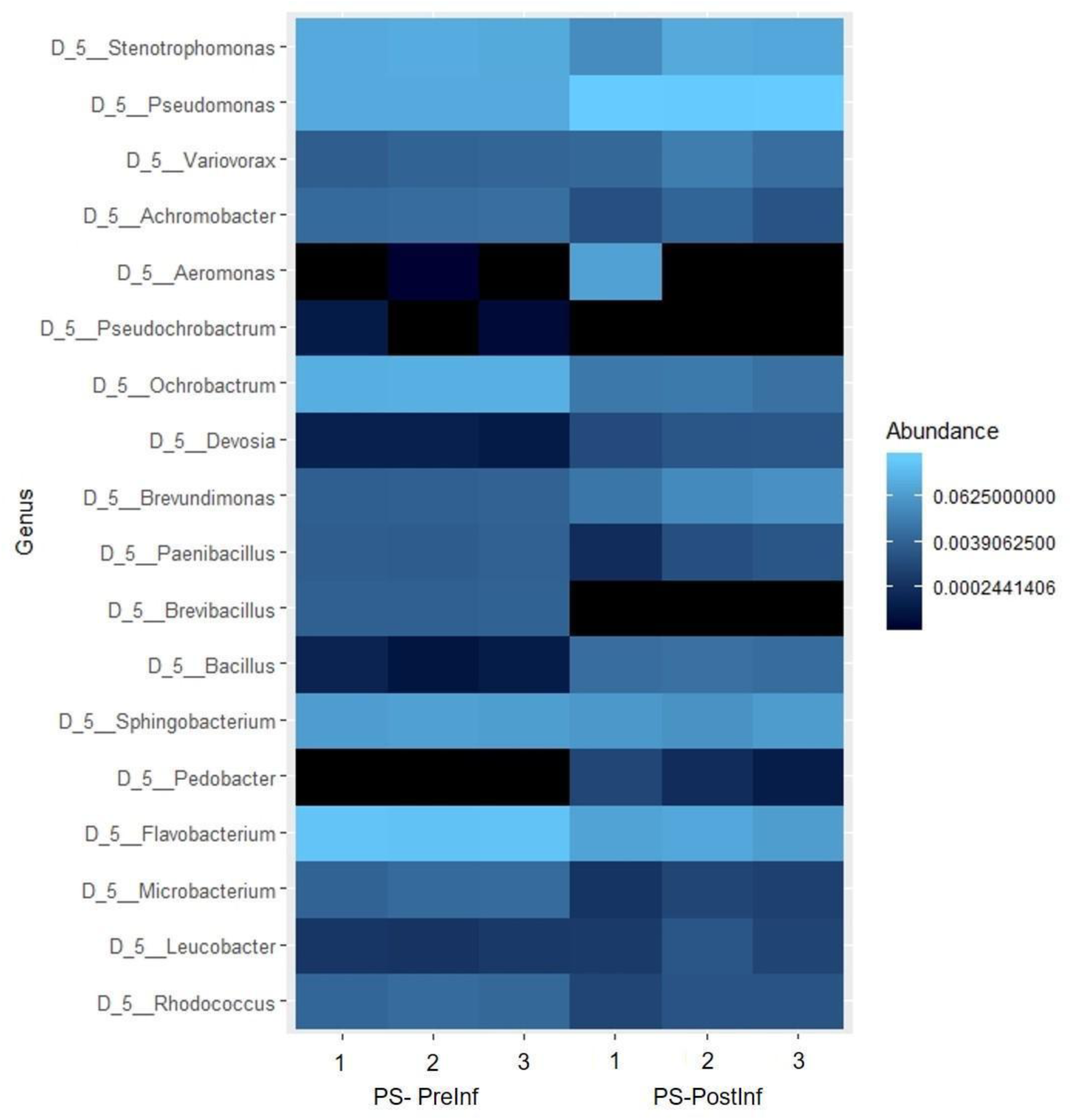
Genus components of microbial communities in each PS biological replicate.

## Discussion

We performed a 16S rRNA-based analysis of the *P. hermaphrodita* microbiota sampled immediately before and after infection of host grey field slugs. We hypothesized that varying microbiomes associated with *P. hermaphrodita* would differentially impact the nematode’s ability to infect slugs, either through direct effects (e.g., direct involvement of bacteria in slug mortality) or through indirect means (e.g., through affecting nematode nutritional metabolism or similar). We observed that slugs exposed to nematodes grown on both concentrations of the *E. coli* treatment survived for significantly longer than slugs exposed to nematodes with the high concentration of both *Pseudomonas* and the original bacterial community. *D. reticulatum* exposed to the low concentration of the original bacterial community survived for significantly fewer days when compared to high *E. coli* rate treatment. These observations support our hypothesis and suggest that the microbiome variation influences the nematode’s ability to infect and kill slugs.

### Bacterial communities associated with P. hermaphrodita

We also hypothesized changes in nematode-associated microbiomes pre-to post-infection samples. The infection of a slug host might act as a selection pressure on the bacterial community, favoring the bacteria that can better contribute to the nematode’s virulence, spread, reproduction, and killing efficacy. Further, recent research has shown differential microbiome responses between nematode-susceptible and nematode-resistant host gastropod species (Sheehy et al. 2023).

The microbiome samples of all the EC, PS, and BC samples showed complex taxa (ASVs) compositions. The species richness and diversity of EC-PreInf and PS-PreInf communities were higher than our expectations, given that the nematodes had been bleached with hypochlorite, washed thoroughly with water and salt solutions, and reared on cultures of *E. coli* or *Pseudomonas spp.* before the infection assay. The intended bacteria were indeed present in the samples collected before infection trials; however, *Pseudomonas sp.* and *E. coli* accounted for a relatively minor component of the community by the time the infection experiments began. There are many likely contributors to the unexpectedly high taxonomic diversity observed in the EC and PS samples, including environmental bacteria in the lab, certain bacteria surviving the bleaching procedures, and the possibility that there are unknown bacterial endosymbionts associated with *P. hermaphrodita* as has been reported in other nematode systems (Brown et al., 2018; Mobasseri et al., 2019).

Before the infection trial, fifteen different phyla and 82 genera were observed in the BC-PreInf samples, with *Pseudochrobactrum, Flavobacterium, Raoultella,* and *Pseudomonas* being the four most abundant taxa in the community. These four genera all have been reported in association or rivalry with nematodes, for example, *Pseudochrobactrum spp*. with the EPN *Steinernema* (Ogier et al., 2020), *Raoultella spp.* with the root-knot and pine wood nematodes (Liu et al., 2019), *Flavobacterium sp.* With the insect-parasitic nematode *Rhabditis blumi* (Woong et al., 2011), the soybean cyst nematode *Heterodera glycines* (Nour et al., 2003), the oriental beetle pathogenic nematode *Butlerius sp*. (Yi et al., 2007), and *Pseudomonas* spp. with *C. elegans* (Pees et al., 2024). The high-abundance components in BC-PreInf samples are relatively similar to that of *C. elegans*, whose dominant genera are *Pseudomonas, Stenotrophomonas, Ochrobactrum, Sphingomonas*, and unclassified Enterobacteriaceae (*Raoultella* is a representative of this family) (Dirksen et al., 2016; Zhang et al., 2017). *Stenotrophomonas, Ochrobactrum,* and *Sphingomonas* spp. were all detected in the top 10 most abundant genera in our BC-PreInf bacterial community.

The complex nature of the bacterial community structure and the results of slug mortality during the infection assay indicated that *P. hermaphrodita* could carry out slug infection with various bacteria regardless of the primary bacterial food source, although at different rates. This observation agrees with a previous finding that the nematode is able to associate with non-specific and complex microbial assemblages (Rae et al., 2010). However, the slug-killing efficacy clearly varied between different bacteria-nematode combinations. The BC-cultured nematodes were the first out of the three treatments to cause significant slug mortality compared to the negative controls on Day 5, followed by the PS nematodes on Day 6, and lastly, the EC nematodes, starting Day 10. However, the nematodes treated with *Pseudomonas* took only three days to kill slugs and were the first to cause 100% slug mortality, followed by the BC nematodes (five days) and EC nematodes (ten days). The results may imply that *Pseudomonas* spp. might influence the nematode’s pathogenicity, possibly by assisting *P. hermaphrodita*’s virulence and/or providing a favorable food source, which may benefit the worm’s growth and development for better infection.

Analysis of ASVs in both the PS-PreInf and PS-PostInf sample sets revealed four common ASVs that belonged to the genus *Pseudomonas* (see Results). Out of these four ASVs, we detected the presence of one particular ASV in all sample sets, of which the relative abundance consistently increased after the course of infection in all three bacterial treatments. This suggests that the specific ASV might be stably retained with the nematode and could be important to the slug infection activity, although laboratory contamination cannot be ruled out since it was observed in all treatments. The abundance of three other common *Pseudomonas* ASVs before and after infection also expanded.

### Bacterial community shifts after slug infection

In general, we found that α-diversity indices were higher in all of the post-infection samples as compared to pre-infection samples, especially the *Pseudomonas* and bacterial community treatments. For BC-PostInf samples, Chao1 and Faith’s PD α-diversity indices increased significantly, which may indicate the enrichment of low-abundance taxa during the course of infection and also implied that the diversity in the PostInf microbiome is not just the result of the presence of a few highly diverse taxonomic groups, but rather a broader range of phylogenetically different taxa. Indeed, for example, the total number of different genera in the BC microbiomes more than tripled after infection, however, the composition of the top 20 most abundant taxa remained stable. Likewise, for the *Pseudomonas-*enriched treatment, we reported a significant increase in Chao1 and Shannon indices after the assay, indicating a spike in rare taxa post-infection, although composition analysis pointed to the dominance of taxa of the genus *Pseudomonas* in the community.

### Microbial community differs by type of bacterial treatment and time of collection

We found that the difference between samples from different bacteria-nematode combinations was significant for Bray-Curtis and weighted UniFrac metrics, but not for unweighted UniFrac. This suggests that between bacterial treatments, the communities tend to have a common set of core ASVs (i.e. similar richness), but these ASVs have highly varied abundances (i.e. diversity). Additionally, it is likely that the higher abundance taxa are distinct between samples from different bacterial treatments, with shared taxa of low abundance.

In the meantime, samples collected before and after infection did not show a significant difference by the weighted UniFrac metric, indicating that only low-abundance taxa differ between the communities pre- and post-infection. This observation is in concordance with the findings of the α-diversity analyses discussed above.

## Conclusion

In this study, we assessed the pathogenicity of *P. hermaphrodita* in association with different bacteria and reported the compositional changes in the nematode’s microbiome before and after slug infection. We observed a substantial increase in species richness across treatments; however, the composition of the most abundant taxa pre-infection remained mostly the same as compared to post-infection. We also detected four specific taxa of the genus *Pseudomonas*, whose relative abundance expanded remarkably after infection in the communities originally enriched with *Pseudomonas.* This finding suggests future investigations to further evaluate the possible role of *Pseudomonas sp.* in the killing of slugs.

## References

Adomaitis, M., Skujienė, G., and Račinskas, P. (2022). Reducing the Application Rate of Molluscicide Pellets for the Invasive Spanish Slug, Arion vulgaris. Insects 13. doi: 10.3390/insects13030301

Aktar, M. W., Sengupta, D., and Chowdhury, A. (2009). Impact of pesticides use in agriculture: their benefits and hazards. Interdiscip. Toxicol. 2, 1.

Anderson, N. P., Hoffman, G. D., and Dreves, A. J. (2011). Evaluation of newly formulated molluscides for control of slugs in western Oregon grass seed fields. *Seed Production Research*, Oregon State University 130, 15–18.

Balvočiūtė, M., and Huson, D. H. (2017). SILVA, RDP, Greengenes, NCBI and OTT - how do these taxonomies compare? BMC Genomics 18. doi: 10.1186/s12864-017-3501-4

Barker, G. M. (2002). Molluscs as crop pests. Wallingford, England: CABI Publishing.

Brown, A. M. V., Wasala, S. K., Howe, D. K., Peetz, A. B., Zasada, I. A., and Denver, D. R. (2018). Comparative Genomics of Wolbachia–Cardinium Dual Endosymbiosis in a Plant-Parasitic Nematode. Front. Microbiol. 9, 414591.

Callahan, B. J., McMurdie, P. J., and Holmes, S. P. (2017). Exact sequence variants should replace operational taxonomic units in marker-gene data analysis. ISME J. 11. doi: 10.1038/ismej.2017.119

Callahan, B. J., McMurdie, P. J., Rosen, M. J., Han, A. W., Johnson, A. J., and Holmes, S. P. (2016). DADA2: High-resolution sample inference from Illumina amplicon data. Nat. Methods 13. doi: 10.1038/nmeth.3869

Centers for Disease Control and Prevention (2023). Parasitic Meningitis. cdc.gov. Available at: https://www.cdc.gov/meningitis/parasitic.html (Accessed April 29, 2024).

Chao, A. (1984). Nonparametric Estimation of the Number of Classes in a Population. Scandinavian Journal of Statistics 11, 265–270.

Davis, N. M., Proctor, D. M., Holmes, S. P., Relman, D. A., and Callahan, B. J. (2018). Simple statistical identification and removal of contaminant sequences in marker-gene and metagenomics data. Microbiome 6. doi: 10.1186/s40168-018-0605-2

De Ley, I. T., McDonnell, R. D., Lopez, S., Paine, T. D., and De Ley, P. (2014). Phasmarhabditis hermaphrodita (Nematoda: Rhabditidae), a potential biocontrol agent isolated for the first time from invasive slugs in North America. Nematology 16, 1129–1138.

Denver, D. R., Howe, D. K., Colton, A. J., Richart, C. H., and Mc Donnell, R. J. (2024). The biocontrol nematode *Phasmarhabditis hermaphrodita* infects and increases mortality of *Monadenia fidelis*, a non-target terrestrial gastropod species endemic to the Pacific Northwest of North America, in laboratory conditions. PLoS One 19, e0298165.

Dirksen, P., Marsh, S. A., Braker, I., Heitland, N., Wagner, S., Nakad, R., et al. (2016). The native microbiome of the nematode Caenorhabditis elegans: gateway to a new host-microbiome model. BMC Biol. 14. doi: 10.1186/s12915-016-0258-1

Faith, D. P. (1992). Conservation evaluation and phylogenetic diversity. Biol. Conserv. 61, 1–10.

Forst, S., Dowds, B., Boemare, N., and Stackebrandt, E. (1997). *Xenorhabdus* and *Photorhabdus* spp.: bugs that kill bugs. Annu. Rev. Microbiol. 51, 47–72.

Godan, D. (1983). Pest Slugs and Snails: Biology and Control. Springer Berlin Heidelberg.

Howe, D. K., Ha, A. D., Colton, A., De Ley, I. T., Rae, R. G., Ross, J., et al. (2020). Phylogenetic evidence for the invasion of a commercialized European Phasmarhabditis hermaphrodita lineage into North America and New Zealand. PLoS One 15. doi: 10.1371/journal.pone.0237249

Katoh, K., Misawa, K., Kuma, K., and Miyata, T. (2002). MAFFT: a novel method for rapid multiple sequence alignment based on fast Fourier transform. Nucleic Acids Res. 30. doi: 10.1093/nar/gkf436

Kembel, S. W., Cowan, P. D., Helmus, M. R., Cornwell, W. K., Morlon, H., Ackerly, D. D., et al. (2010). Picante: R tools for integrating phylogenies and ecology. Bioinformatics 26, 1463– 1464.

Klindworth, A., Pruesse, E., Schweer, T., Peplies, J., Quast, C., Horn, M., et al. (2013). Evaluation of general 16S ribosomal RNA gene PCR primers for classical and next-generation sequencing-based diversity studies. Nucleic Acids Res. 41. doi: 10.1093/nar/gks808

Lacey, L. A., and Georgis, R. (2012). Entomopathogenic nematodes for control of insect pests above and below ground with comments on commercial production. J. Nematol. 44, 218– 225.

Liu, Y., Ponpandian, L. N., Kim, H., Jeon, J., Hwang, B. S., Lee, S. K., et al. (2019). Distribution and diversity of bacterial endophytes from four Pinus species and their efficacy as biocontrol agents for devastating pine wood nematodes. Sci. Rep. 9, 1–12.

Mc Donnell, R. J., and Anderson, N. P. (2017). Pest Slugs in Western Oregon Seed Crops: Stakeholder Knowledge, Baiting Strategies, and Attitude Toward Novel Management Tools. in Seed Production Research Report (Oregon State University), 1–4.

Mc Donnell, R. J., Lutz, M. S., Howe, D. K., and Denver, D. R. (2018). First Report of the Gastropod-Killing Nematode, Phasmarhabditis hermaphrodita, in Oregon, U.S.A. J. Nematol. 50, 77.

Mc Donnell, R. J., Paine, T. D., and Gormally, M. J. (2009). Slugs: A Guide to the Invasive and Native Fauna of California. UCANR Publications.

Mc Donnell, R.J., Colton, A.J., Howe, D.K., and Denver, D.R. (2020). Lethality of four species of *Phasmarhabditis* (Nematoda: Rhabditidae) to the invasive slug, *Deroceras reticulatum* (Gastropoda: Agriolimacidae) in laboratory infectivity trials. Biol Control 150: 104349.

McMurdie, P. J., and Holmes, S. (2013). phyloseq: An R Package for Reproducible Interactive Analysis and Graphics of Microbiome Census Data. PLoS One 8, e61217.

Mobasseri, M., Hutchinson, M. C., Afshar, F. J., and Pedram, M. (2019). New evidence of nematode-endosymbiont bacteria coevolution based on one new and one known dagger nematode species of Xiphinema americanum-group (Nematoda, Longidoridae). PLoS One 14, e0217506.

Nour, S. M., Lawrence, J. R., Zhu, H., Swerhone, G. D. W., Welsh, M., Welacky, T. W., et al. (2003). Bacteria Associated with Cysts of the Soybean Cyst Nematode (Heterodera glycines). Appl. Environ. Microbiol. doi: 10.1128/AEM.69.1.607-615.2003

Ogier, J.-C., Pagès, S., Frayssinet, M., and Gaudriault, S. (2020). Entomopathogenic nematode-associated microbiota: from monoxenic paradigm to pathobiome. Microbiome 8, 1–17.

Oksanen, J., Guillaume Blanchet, F., Kindt, R., Legendre, P., Minchin, P. R., O’Hara, R. B., et al. (2012). vegan: Community Ecology Package. Available at: https://researchportal.helsinki.fi/en/publications/vegan-community-ecology-package (Accessed April 30, 2024).

Pees, B., Peters, L., Treitz, C., Hamerich, I. K., Kissoyan, K. A. B., Tholey, A., et al. (2024). The Caenorhabditis elegans proteome response to two protective Pseudomonas symbionts. MBio. doi: 10.1128/mbio.03463-23

Price, M. N., Dehal, P. S., and Arkin, A. P. (2010). FastTree 2--approximately maximum-likelihood trees for large alignments. PLoS One 5. doi: 10.1371/journal.pone.0009490

Quast, C., Pruesse, E., Yilmaz, P., Gerken, J., Schweer, T., Yarza, P., et al. (2013). The SILVA ribosomal RNA gene database project: improved data processing and web-based tools. Nucleic Acids Res. 41, D590–D596.

Rae, R. G., Tourna, M., and Wilson, M. J. (2010). The slug parasitic nematode Phasmarhabditis hermaphrodita associates with complex and variable bacterial assemblages that do not affect its virulence. J. Invertebr. Pathol. 104. doi: 10.1016/j.jip.2010.04.008

Rae, R., Verdun, C., Grewal, P. S., Robertson, J. F., and Wilson, M. J. (2007). Biological control of terrestrial molluscs using Phasmarhabditis hermaphrodita--progress and prospects. Pest Manag. Sci. 63, 1153–1164.

Sheehy, L., Cutler, J., Weedall, G. D., and Rae, R. (2022). Microbiome Analysis of Malacopathogenic Nematodes Suggests No Evidence of a Single Bacterial Symbiont Responsible for Gastropod Mortality. Front. Immunol. 13, 878783.

Sicard, M., Ferdy, J.-B., Pagès, S., Le Brun, N., Godelle, B., Boemare, N., et al. (2004). When mutualists are pathogens: an experimental study of the symbioses between Steinernema (entomopathogenic nematodes) and Xenorhabdus (bacteria). J. Evol. Biol. 17, 985–993.

Simpson, E. H. (1949). Measurement of Diversity. Nature 163, 688–688.

Spellerberg, I. F., and Fedor, P. J. (2003). A tribute to Claude Shannon (1916–2001) and a plea for more rigorous use of species richness, species diversity and the “Shannon–Wiener” Index. Glob. Ecol. Biogeogr. 12, 177–179.

Stiernagle, T. (2006). Maintenance of C. elegans. WormBook. doi: 10.1895/wormbook.1.101.1

Tan, L., and Grewal, P. S. (2001a). Infection behavior of the rhabditid nematode Phasmarhabditis hermaphrodita to the grey garden slug Deroceras reticulatum. J. Parasitol. 87. doi: 10.1645/0022-3395(2001)087[1349:IBOTRN]2.0.CO;2

Tan, L., and Grewal, P. S. (2001b). Pathogenicity of Moraxella osloensis, a Bacterium Associated with the Nematode Phasmarhabditis hermaphrodita, to the Slug Deroceras reticulatum. Appl. Environ. Microbiol. 67, 5010.

Tan, L., and Grewal, P. S. (2002). Endotoxin activity of Moraxella osloensis against the grey garden slug, Deroceras reticulatum. Appl. Environ. Microbiol. 68. doi: 10.1128/AEM.68.8.3943-3947.2002

Tudi, M., Ruan, H. D., Wang, L., Lyu, J., Sadler, R., Connell, D., et al. (2021). Agriculture Development, Pesticide Application and Its Impact on the Environment. Int. J. Environ. Res. Public Health 18. doi: 10.3390/ijerph18031112

UK Department for Environment (2020). Outdoor use of metaldehyde to be banned to protect wildlife. GOV.UK. Available at: https://www.gov.uk/government/news/outdoor-use-of-metaldehyde-to-be-banned-to-protect-wildlife (Accessed April 29, 2024).

Wilson, M. J., Glen, D. M., and George, S. K. (1993). The rhabditid nematode Phasmarhabditis hermaphrodita as a potential biological control agent for slugs. Biocontrol Sci. Technol. 3, 503–511.

Wilson, M. J., Glen, D. M., George, S. K., and Pearce, J. D. (1995a). Monoxenic culture of the slug parasite Phasmarhabditis hermaphrodita (Nematoda: Rhabditidae) with different bacteria in liquid and solid phase. Fundam Appl Nematol 2, 159–166.

Wilson, M. J., Glen, D. M., George, S. K., and Pearce, J. D. (1995b). Selection of a bacterium for the mass production of Phasmarhabditis hermaphrodita (Nematoda: Rhabditidae) as a biocontrol agent for slugs. Fundamental and Applied Nematology 5, 419–425.

Wilson, M. J., Glen, D. M., Pearce, J. D., and Rodgers, P. B. (1995c). Monoxenic culture of the slug parasite Phasmarhabditis hermaphrodita (Nematoda: Rhabditidae) with different bacteria in liquid and solid phase-fdi:42721-Horizon. Fundamental and Applied Nematology 2, 159–166.

Woong, P., Ook, K., HaJae-Seok, Hun, Y., Hwan, K., BilgramiAnwar, L., et al. (2011). Effects of associated bacteria on the pathogenicity and reproduction of the insect-parasitic nematode Rhabditis blumi (Nematoda: Rhabditida). Can. J. Microbiol. doi: 10.1139/w11-067

Yi, Y. K., Park, H. W., Shrestha, S., Seo, J., Kim, Y. O., Shin, C. S., et al. (2007). Identification of two entomopathogenic bacteria from a nematode pathogenic to the Oriental beetle, Blitopertha orientalis. J. Microbiol. Biotechnol. 17. Available at: https://pubmed.ncbi.nlm.nih.gov/18050915/ (Accessed April 30, 2024).

Zhang, F., Berg, M., Dierking, K., Félix, M. A., Shapira, M., Samuel, B. S., et al. (2017). Caenorhabditis elegans as a Model for Microbiome Research. Front. Microbiol. 8. doi: 10.3389/fmicb.2017.00485

