## Supplemental Materials for "Microbiome dynamics associated with the infection of grey field slugs by the biocontrol nematode *Phasmarhabditis hermaphrodita*"

### Supplementary material

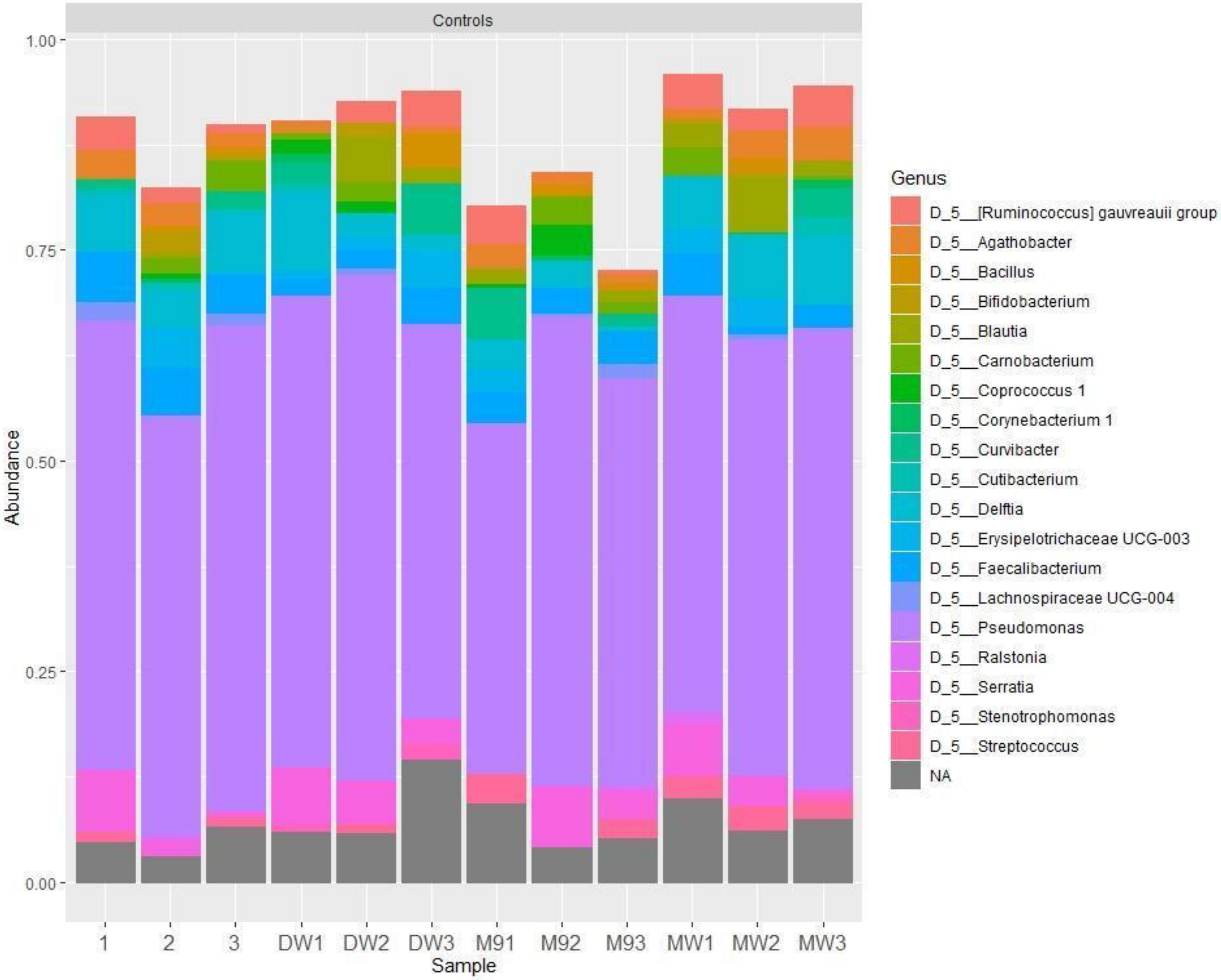


Figure S1. Composition of ASVs contaminants removed from ASV table.

Table S1. Naming scheme of samples

| **Bacterial treatment** | **Time point** | **Sample name** |
|---|---|---|
| *E. coli*-enriched samples | before infection | EC-PreInf |
| | after infection | EC-PostInf |
| *Pseudomonas* sp. - enriched samples | before infection | PS-PreInf |
| | after infection | PS-PostInf |
| Complex bacterial community | before infection | BC-PreInf |
| | after infection | BC-PostInf |

Table S2. The number of quality-controlled read counts in decontaminated samples.

| Sample | No. of reads | Total reads in each combination | Median value in each combination | Mean value in each combination |
|---|---|---|---|---|
| EA1 | 604,813 | 4,087,108 | 490,705 | 681,185 |
| EA2 | 1,610,879 | | | |
| EA3 | 1,091,611 | | | |
| EB1 | 224,670 | | | |
| EB2 | 376,596 | | | |
| EB3 | 178,539 | | | |
| UA1 | 153,427 | 2,572,446 | 231,484 | 428,741 |
| UA2 | 153,680 | | | |
| UA3 | 100,672 | | | |
| UB1 | 309,288 | | | |

| | | | | |
|---|---|---|---|---|
| UB2 | 1,005,733 | | | |
| UB3 | 849,646 | | | |
| PA1 | 641,501 | 3,762,641 | 453,176 | 627,107 |
| PA2 | 1,198,645 | | | |
| PA3 | 1,231,796 | | | |
| PB1 | 264,851 | | | |
| PB2 | 250,356 | | | |
| PB3 | 175,492 | | | |

Table S3: **(A)**. Phylum composition and relative abundance of the bacterial community associated with *P. hermaphrodita* before infection (samples BC-PreInf). Phyla were sorted from the highest to lowest mean abundance in samples.

| Phylum | meanRA | sdRA | minRA | maxRA |
|---|---|---|---|---|
| D_1__Bacteroidetes | 0.022711 | 0.05372 | 1.30E-05 | 0.216386 |
| D_1__Proteobacteria | 0.0099 | 0.025887 | 1.99E-05 | 0.195276 |
| D_1__Gemmatimonadetes | 0.002883 | 0.004853 | 7.81E-05 | 0.010142 |
| D_1__Firmicutes | 0.002073 | 0.003461 | 1.95E-05 | 0.012119 |
| D_1__Chloroflexi | 0.001788 | NA | 0.001788 | 0.001788 |
| D_1__Planctomycetes | 0.001664 | 0.001337 | 3.90E-05 | 0.003268 |
| D_1__Acidobacteria | 0.001592 | 0.000449 | 0.001222 | 0.002245 |
| NA | 0.001585 | 0.002498 | 1.30E-05 | 0.006981 |
| D_1__Actinobacteria | 0.001375 | 0.001653 | 3.26E-05 | 0.006019 |
| D_1__Verrucomicrobia | 0.001323 | 0.001488 | 0.000169 | 0.003447 |

| | | | | |
|---|---|---|---|---|
| D_1__WS2 | 0.000864 | NA | 0.000864 | 0.000864 |
| D_1__Armatimonadetes | 0.000808 | NA | 0.000808 | 0.000808 |
| D_1__Patescibacteria | 0.000505 | 2.30E-05 | 0.000489 | 0.000521 |
| D_1__Cyanobacteria | 0.000192 | 0.000169 | 5.96E-05 | 0.000437 |
| D_1__Deinococcus-Thermus | 4.97E-05 | NA | 4.97E-05 | 4.97E-05 |

**B.** Genus composition of the natural bacterial community associated with *P. hermaphrodita.* Genera were sorted from the highest to lowest mean abundance in samples.

| Genus | meanRA | sdRA | minRA | maxRA |
|---|---|---|---|---|
| D_5__Pseudochrobactrum | 0.069449 | 0.085976 | 0.002215 | 0.195276 |
| D_5__Flavobacterium | 0.045221 | 0.077108 | 1.30E-05 | 0.216386 |
| D_5__Raoultella | 0.01917 | 0.020054 | 0.000404 | 0.062018 |
| D_5__Pseudomonas | 0.018958 | 0.027937 | 4.55E-05 | 0.110072 |
| D_5__Brevundimonas | 0.010851 | 0.011081 | 0.000467 | 0.029283 |
| D_5__Sphingobacterium | 0.007877 | 0.00789 | 0.000235 | 0.026679 |
| D_5__Pedobacter | 0.005992 | 0.002071 | 0.003795 | 0.009097 |
| D_5__wb1-P19 | 0.005553 | NA | 0.005553 | 0.005553 |
| D_5__Ochrobactrum | 0.005453 | 0.003768 | 0.001662 | 0.011023 |
| D_5__Bauldia | 0.005066 | NA | 0.005066 | 0.005066 |
| D_5__Paenibacillus | 0.004858 | 0.004821 | 0.000469 | 0.012119 |
| D_5__Achromobacter | 0.004599 | 0.002172 | 0.001597 | 0.00691 |
| D_5__Kaistia | 0.004247 | 0.004221 | 0.00058 | 0.00886 |
| NA | 0.003932 | 0.007775 | 1.30E-05 | 0.033866 |
| D_5__Sphingomonas | 0.003857 | 0.003819 | 0.000111 | 0.009262 |
| D_5__ADurb.Bin063-1 | 0.003447 | NA | 0.003447 | 0.003447 |
| D_5__Microbacterium | 0.003383 | 0.002822 | 0.000372 | 0.006019 |
| D_5__Rhodococcus | 0.003253 | 0.002015 | 0.000111 | 0.00509 |
| D_5__Stenotrophomonas | 0.003141 | 0.004696 | 1.99E-05 | 0.012148 |
| D_5__Allorhizobium-Neorhizobium-Pararhizobium-Rhizobium | 0.003072 | 0.001925 | 0.000891 | 0.005275 |

| | | | | |
|---|---|---|---|---|
| D_5__Rhodoferax | 0.002959 | NA | 0.002959 | 0.002959 |
| D_5__uncultured | 0.002167 | 0.002845 | 3.90E-05 | 0.010142 |
| D_5__Comamonas | 0.002157 | 0.001104 | 0.001356 | 0.003416 |
| D_5__Anaerococcus | 0.002007 | NA | 0.002007 | 0.002007 |
| D_5__Bacteroides | 0.001973 | 0.002716 | 5.21E-05 | 0.003894 |
| D_5__Marmoricola | 0.001828 | 0.001888 | 0.000104 | 0.00449 |
| D_5__uncultured soil bacterium | 0.001788 | NA | 0.001788 | 0.001788 |
| D_5__Herminiimonas | 0.001429 | 0.000767 | 0.000886 | 0.001972 |
| D_5__Chryseobacterium | 0.001421 | NA | 0.001421 | 0.001421 |
| D_5__Caulobacter | 0.001351 | 0.001301 | 5.96E-05 | 0.003084 |
| D_5__Propionivibrio | 0.001297 | NA | 0.001297 | 0.001297 |
| Ambiguous_taxa | 0.001281 | 0.000945 | 0.000613 | 0.001949 |
| D_5__Oikopleura dioica | 0.001245 | NA | 0.001245 | 0.001245 |
| D_5__RB41 | 0.001222 | NA | 0.001222 | 0.001222 |
| D_5__Porphyrobacter | 0.001162 | NA | 0.001162 | 0.001162 |
| D_5__uncultured bacterium | 0.001155 | 0.001066 | 0.000489 | 0.003258 |
| D_5__Terrimonas | 0.001108 | NA | 0.001108 | 0.001108 |
| D_5__Nakamurella | 0.001108 | NA | 0.001108 | 0.001108 |
| D_5__Nesterenkonia | 0.001088 | NA | 0.001088 | 0.001088 |
| D_5__Rathayibacter | 0.001036 | NA | 0.001036 | 0.001036 |
| D_5__Crossiella | 0.000939 | NA | 0.000939 | 0.000939 |
| D_5__Leucobacter | 0.000917 | 0.000763 | 0.00013 | 0.002781 |
| D_5__Lachnoclostridium | 0.000867 | NA | 0.000867 | 0.000867 |
| D_5__Pir4 lineage | 0.00072 | 0.000963 | 3.90E-05 | 0.001401 |

| | | | | |
|---|---|---|---|---|
| D_5__Variovorax | 0.000719 | 0.000572 | 0.000137 | 0.001623 |
| D_5__Friedmanniella | 0.000704 | NA | 0.000704 | 0.000704 |
| D_5__Nordella | 0.000691 | NA | 0.000691 | 0.000691 |
| D_5__Devosia | 0.000677 | 0.000596 | 0.000104 | 0.001744 |
| D_5__Brevibacillus | 0.00062 | 0.000738 | 9.78E-05 | 0.001464 |
| D_5__Phreatobacter | 0.000602 | 0.000382 | 0.000332 | 0.000872 |
| D_5__Shinella | 0.000591 | 0.000626 | 0.000119 | 0.001301 |
| D_5__Faecalibacterium | 0.00055 | 0.000578 | 0.00015 | 0.001212 |
| D_5__Nocardioides | 0.000521 | 0.000516 | 0.000156 | 0.000886 |
| D_5__Hydrogenophaga | 0.000504 | 0.000621 | 6.52E-05 | 0.000944 |
| D_5__Medicago truncatula | 0.000437 | NA | 0.000437 | 0.000437 |
| D_5__Arenimonas | 0.00043 | 0.000378 | 0.000163 | 0.000697 |
| D_5__Candidatus Udaeobacter | 0.00043 | NA | 0.00043 | 0.00043 |
| D_5__Geodermatophilus | 0.000398 | NA | 0.000398 | 0.000398 |
| D_5__Gemmatimonas | 0.000365 | NA | 0.000365 | 0.000365 |
| D_5__Streptococcus | 0.000328 | NA | 0.000328 | 0.000328 |
| D_5__Jatrophihabitans | 0.000274 | NA | 0.000274 | 0.000274 |
| D_5__Moheibacter | 0.00026 | NA | 0.00026 | 0.00026 |
| D_5__Roseburia | 0.000254 | 0.000323 | 2.60E-05 | 0.000482 |
| D_5__Sphingopyxis | 0.000247 | 0.00023 | 8.47E-05 | 0.00041 |
| D_5__uncultured Sphingomonadaceae bacterium | 0.000235 | NA | 0.000235 | 0.000235 |
| D_5__alphaI cluster | 0.000235 | NA | 0.000235 | 0.000235 |
| D_5__Coprococcus 1 | 0.000215 | NA | 0.000215 | 0.000215 |
| D_5__Gaiella | 0.000202 | NA | 0.000202 | 0.000202 |

| | | | | |
|---|---|---|---|---|
| D_5__Leptothrix | 0.000182 | NA | 0.000182 | 0.000182 |
| D_5__Prosthecobacter | 0.000169 | NA | 0.000169 | 0.000169 |
| D_5__Aeromicrobium | 0.000163 | NA | 0.000163 | 0.000163 |
| D_5__Rubellimicrobium | 0.00015 | NA | 0.00015 | 0.00015 |
| D_5__Candidimonas | 0.000143 | NA | 0.000143 | 0.000143 |
| D_5__Mesorhizobium | 0.00014 | 0.000124 | 5.21E-05 | 0.000228 |
| D_5__Delftia | 7.81E-05 | NA | 7.81E-05 | 7.81E-05 |
| D_5__metagenome | 7.81E-05 | NA | 7.81E-05 | 7.81E-05 |
| D_5__Rhodopseudomonas | 6.52E-05 | NA | 6.52E-05 | 6.52E-05 |
| D_5__Deinococcus | 4.97E-05 | NA | 4.97E-05 | 4.97E-05 |
| D_5__Mycobacterium | 3.91E-05 | NA | 3.91E-05 | 3.91E-05 |
| D_5__Conexibacter | 3.26E-05 | NA | 3.26E-05 | 3.26E-05 |
| D_5__Hyphomicrobium | 2.60E-05 | NA | 2.60E-05 | 2.60E-05 |
| D_5__Dorea | 1.95E-05 | NA | 1.95E-05 | 1.95E-05 |
